# Causal Inference in Microbiomes Using Intervention Calculus

**DOI:** 10.1101/2020.02.28.970624

**Authors:** Musfiqur Rahman Sazal, Vitalii Stebliankin, Kalai Mathee, Giri Narasimhan

## Abstract

Inferring causal effects is critically important in biomedical research as it allows us to move from the typical paradigm of associational studies to causal inference, and can impact treatments and therapeutics. Association patterns can be coincidental and may lead to wrong inferences in complex systems. Microbiomes are highly complex, diverse, and dynamic environments. Microbes are key players in health and diseases. Hence knowledge of genuine causal relationships among the entities in a microbiome, and the impact of internal and external factors on microbial abundance and interactions are essential for understanding disease mechanisms and making treatment recommendations.

In this paper, we investigate fundamental causal inference techniques to measure the causal effects of various entities in a microbiome. In particular, we show how to use these techniques on microbiome datasets to study the rise and impact of antibiotic-resistance in microbiomes. Our main contributions include the following. We introduce a novel pipeline for microbiome studies, new ideas for experimental design under weaker assumptions, and data augmentation by context embedding. Our pipeline is robust, different from traditional approaches, and able to predict interventional effects without any controlled experiments. Our work shows the advantages of causal inference in identifying potential pathogenic, beneficial, and antibiotic-resistant bacteria. We validate our results using results that were previously published.

## 1 Introduction

Inferring causality is the process of connecting a cause with an effect. Identifying even a single causal relationship from data is often worth more than observing dozens of correlations in a data set. The study of causality is not new in many areas of science, but in recent years with advances in causal calculus, data science, and machine learning, the focus is now on how to draw causal conclusions in a data-driven way. Given a sufficiently large and rich data set, the theoretical foundations of causality allows us to go beyond merely discovering statistical associations in data, to infer quantitative causal relationships and to explore “what-if” questions, thus profoundly impacting data-driven decision making in many domains. In the field of biomedicine, inferring causal relationships could impact treatment and therapy.

Causal inference can be achieved if a causal structure is readily available based on prior knowledge from experts (e.g., exercise reduces cholesterol). However, in most real-world situations, this is not known. Alternatively, causal inference is also possible if extensive experimentation is possible along with the ability to control all variables in play. In most applications, controlling all variables is impossible (e.g., setting the abundance of bacteria *A* or the concentration of metabolite *M* in a person’s gut to specific values). If both options are unavailable, but extensive observational data is available, then we rely on the fact that we can test whether a causal model fits the data, even though no experimental manipulation has been carried out. Artificial intelligence already showed huge success in many domains, for example, computer vision [60, 22, 46] and speech recognition [20]. However, causal inference has been compared to human level intelligence [38] and recently has been successfully applied to data from education [38], economics [54], online advertising [5], medicine and epidemiology [25, 9], social sciences [37], natural language processing [59], policy evaluation [51], recommendation systems [4], and much more.

A *microbiome* is a community of microbes including bacteria, archaea, protists, fungi and viruses that share an environmental niche [41]. Microbiomes have been referred to as a *social network* because of the complex set of potential interactions between its members [15, 14], their products and their host. These interactions take the form of cellular communications, cooperation, competition, and much more. All animals and plants live in close association with communities of microbes [30]. These niche communities are highly dynamic. Human bodies harbor rich communities of microbes mostly in the gastrointestinal and reproductive tracts, and on cutaneous and mucosal surfaces such as the skin and the oral cavity [11, 40]. Bacteria (and microbes, in general) play an essential role in human health by helping in a variety of routine processes including digestion, immune responses, and synthesis of useful vitamins and other metabolites. Interactions between microbes in these communities can impact the genes they express and the metabolites they produce or utilize, and can therefore impact the health of the host or the environmental niche [13]. In a *symbiotic* microbiome, many microbial taxa play a useful role leading to a healthy ecosystem. An imbalance (dysbiosis) in the microbial community is strongly associated with a variety of human diseases [34], often by producing harmful metabolites or by preventing the production of sufficient quantities of necessary products [2]. Thus, inferring causal relationships among the entities of a microbiome and with their hosts are crucial for selection of treatments and recommendation of probiotics [7].

In this paper we investigate the causal relationships between microbes and other entities of the microbiome in subjects with *Inflammatory Bowel Disease* (IBD), with special emphasis on the causal effects of different antibiotics and on the resulting rise of antibiotic resistance in different taxa in the microbiome.

Dysbiosis of the gut microbiome is associated with IBD, colorectal cancer, obesity, and much more. However, the relationships between microbial taxa are complex and the experiments required to understand the causal mechanisms are expensive and time-consuming, and therefore remain poorly understood. Another major threat to public health is the rise of *antibiotic resistance* [56], resulting from the overuse, misuse and abuse of antibiotics. While there is no denying the value of antibiotic treatments to combat infectious diseases [3], the need to study antibiotic resistance as a microbial community characteristic is well recognized as a high priority [21].

Causality in microbiomes is a recent topic of research interest. Bourrat and Fishbach et al. discussed broad ideas about causal inference in microbiomes [6, 16]. Sanna et al. studied causality in microbiome using bidirectional Mendelian randomization [45]. Sazal et al. showed how to extract directional relationships among the taxa from oral microbiomes [48, 47]. Ramakrishnan et al. studied causal relationships in microbiomes related to upper airway diseases [42]. This paper approaches causality in microbiomes using a data-driven approach, drawing whenever possible from appropriate knowledgebases. The only other data-driven approach we found on microbiomes was the work of Mainali et al. [27], where the authors focused on Granger causality, which infers causality from time series data.

## 2 Causal Inference

The first step in inferring causality is to learn the *causal relationships* (also called causal discovery or causal search), which entails discovering the structure of the relationships. The second step is to use the structure to infer the *causal effects*, i.e., the magnitude of the strength of causal relationships.

### 2.1 Causal Discovery

The goal of causal discovery is to establish causal relationships between the entities from observed data or using domain knowledge. A particular type of Bayesian network (BN) is often used to encode such relationships. A BN, sometimes called a belief network or causal network, is a *Probabilistic Graphical Models* (PGMs) that represents a set of variables and their conditional dependencies via a directed acyclic graph (DAG). A causal network is a BN where the edges correspond to direct causal relationships. In a causal network or causal BN, the parents of each vertex are its presumed direct causes. The direct (and indirect) causes of *X_i_* are the variables that, when varied, will change the distribution of *X_i_* [35].

Formally, we define *causal structures* (CS) (or *causal Bayesian networks*) as a class of *PGMs* [36, 24] where each node represents one of *n* random variable from a set, **X** = {*X*_*i*_, *i* = 1,…, *n*}, and each edge represents a direct causal relationship. These structures are represented as a graph *G* = (*V, E*), where each vertex in *V* represents a random variable from **X**, and *E* is the set of edges. Although undirected edges are used in cases where the direction cannot be reliably determined or when both directions appear to be valid, the graph *G* is often “manipulated” to be a Directed Acyclic Graph (DAG). Each random variable *X_i_* has an associated probability distribution. A directed edge in *E* between two vertices represents direct stochastic dependencies. Therefore, if there is no edge connecting two vertices, the corresponding variables are either marginally independent or conditionally independent (conditional on the rest of the variables, or some subset thereof). The “local” probability distribution of a variable *X_i_* depends only on itself and its parents (i.e., the vertices with directed edges into the node *X_i_*); the “global” probability distribution, *P* (**X**) is the product of all local probabilities, i.e., a joint distribution [49], given by

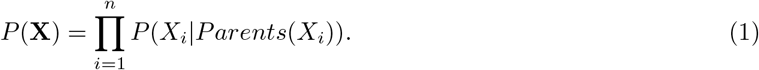

Note that the equation is simpler when the causal structure is sparser. Thus, an important step in our pipeline is to identify all independent pairs of random variables. More importantly, we also identify as many conditionally independent pairs as possible since these represent indirect or non-causal relationships.

All local structures in a causal structure can be classified into three sub-categories: *chains*, *forks*, and *colliders*. In a chain, two variables *X* and *Y* are conditionally independent given *Z*, if there is only one unidirectional path between *X* and *Y*, and *Z* is the set of variables that intercepts that path. In a fork, variable *Z* is a “common cause” for variables *X* and *Y* ; this happens when there is no directed path between *X* and *Y*, and they are independent conditional on *Z*. Finally, variable *Z* is a “collider” node between *X* and *Y*, if it is the “common-effect”. In a collider, as in the fork, there is no directed path between *X* and *Y*. However, the difference is that *X* and *Y* are unconditionally independent, but become dependent when conditioned on *Z* and any descendants of *Z*.

In general, causal models can be very complex. A pair of variables can be connected through multiple chains, forks, and colliders, making it non-trivial to determine the dependency between two arbitrary variables. *Directional separation* (or, just *d-separation*) is a useful concept in this context [18] because covariance terms corresponding to *d*-separated variables are equal to 0. In a directed graph, *G*, two vertices *x* and *y* are *d*-connected if and only if *G* has a collider-free path connecting *x* and *y*. More generally, if *X, Y* and *Z* are disjoint sets of vertices, then *X* and *Y* are *d*-connected by *Z* if and only if *G* has a path *P* between some vertex in *X* and some vertex in *Y* such that for every collider *C* on *P*, either *C* or a descendant *C* is in *Z*, and no non-collider on *P* is in *Z*. *X* and *Y* are *d*-separated by *Z* in *G* if and only if they are not *d*-connected by *Z* in *G*. The concept of *d*-separation allows for more edges to be eliminated in a causal structure.

### 2.2 Intervention

Intervention measures the impact of an action and can be thought of as the effect of “doing/intervening.” It helps to answer interventional questions of the type: “if a person consumes a specific antibiotic, how will the abundance of taxon *A* in her gut change?” or “what is the expected abundance of *B. longum* if the relative abundance of *C. difficile* is fixed at 0.1?” Note that a controlled experiment can potentially answer such interventional questions, but may be either prohibitively expensive, impossible, or unethical to perform. Causal calculus allows us to answer such interventional questions in an *in silico* manner. We clarify that data collected from research studies (e.g., a microbiome study) are observational data, and not the result of controlled interventions, which require that variables be artificially held at specific values. Conditional expectation is given by *E*[*Y*|*X* = *x*], while intervention is given by *E*[*Y*|do(*X* = *x*)], which is the expectation of *Y* if every sample in the population had variable *X* fixed at value *x*. Observational distribution *P* (*y*|*x*) is different from interventional distribution *P* (*y*|do(*x*)). Observational distribution describes that the distribution of *Y* given that variable *X* takes value *x* is observed. On the other hand, interventional distribution of *Y* is what we would observe if we intervened in the data generating process by artificially forcing the variable *X* to take value *x*, but data of other variables remain same. Pearl showed how to compute interventions in a causal model [39]. This is done by “mutilating” the model – to achieve do(*X* = *x*), delete all incoming edges to node *X*, fix its value at *x*, and then perform computations on the resulting network.

### 2.3 Intervention Calculus

Consider the *n* random variables *X*_1_,…, *X*_*n*_ and let *pa_j_* denote the parents of *X_j_*. Any distribution that is generated from a causal structure can be factorized as

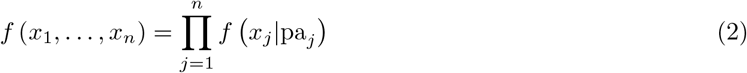

A distribution generated from a DAG with independent error terms results in a Markovian model for which an intervention *do*(*X_i_* = *x*) on the set of variables *X_i_*,…, *X*_*n*_ is given by the following formula

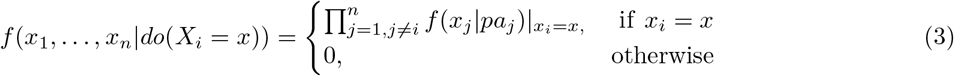

where *f* (*x_j_*|*pa_j_*) are pre-intervention conditional distribution. The above formula uses the causal structure to write interventional distribution on the left-hand side in terms of pre-intervention conditional distributions on the right hand side. It is possible to summarize the distribution generated by an intervention by its mean

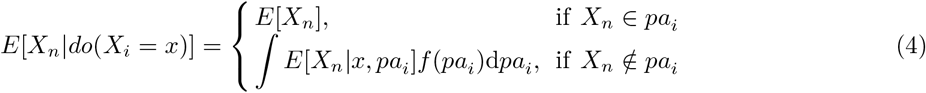

Assuming that the joint distribution of *n* random variables is Gaussian, the causal effect of *X_i_* on *X_n_* is given as

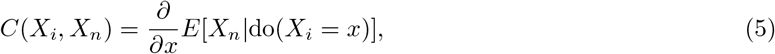

and *γ* becomes constant because of linearity assumption. Since normality implies that *E*(*X_n_*|*pa*_*i*_, *X*_*i*_ = *x*) is linear in *x* and *pa_i_* we can express the expectation value using the following equation

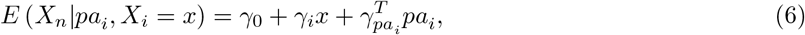

for some values *γ*_0_, *γ_i_* ∈ **R**. The causal effect of *X_i_* on *X_n_* with *X_n_* ∉ *pa_i_* is denoted by *C*(*X_i_, X_n_*) and equals the regression coefficient of *x_i_* above. Thus,

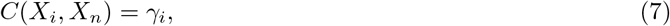

where *γ_i_* is as dictated by Eq. (6).

## 3 Methods and Experiments

In our novel approach, we applied intervention calculus or do-calculus to the causal network constructed from microbiome data sets to (a) determine the causal effects of each microbial taxa on other microbial taxa, and (b) to determine the effects of antibiotics on different microbial taxa. Our proposed method is as follows: (1) learn a causal graph, (2) compute causal effects, (3) analyze the role of causally significant microbial taxa.

### 3.1 Problem Formulation

The problem formulation for causal effects among taxa is as follows: Let *T* = {*B*_1_, *B*_2_,…, *B_n_*} be the set of microbial taxa present in the cohort of healthy or disease samples with abundance values {*b*_1_, *b*_2_,…, *b_n_*}. For two taxa {*B_i_, B_j_*} ∈ *T*, the causal effect of *B_i_* on *B_j_* is given by

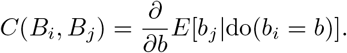

We computed causal effects for all pairs in *T* and ranked all taxa according to the sum of absolute values of causal effect on all the other taxa, with the hope of identifying the most influential taxa in the microbiome. Using the above ranking, we considered the top 30% of taxa for further analysis.

Similarly, to study the causal effects of different antibiotics on the taxa, we let *T* = {*A*_1_, *A*_2_,…, *A_n_*} be the set of antibiotics applied, and let *O* = {*b*_1_, *b*_2_,…, *b*_*n*_} be the set of abundances of the microbial taxa {*B*_1_, *B*_2_,…, *B_n_*} for those samples. (*T* is for treatment or interventional variable, *O* is for outcome variable in this context.) Causal effect of an antibiotic *A_i_* on the abundance of a microbial taxon *b_i_* is 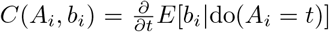. Since causal effects can be positive or negative, we computed causal effects for all pairs in *T* × *O* and separately ranked the pairs with positive and negative causal effects for further analysis.

### 3.2 Data

We analyzed five data sets related to IBD: three from Integrative Human Microbiome Project (iHMP) [1] and two from MicrobiomeHD database [12]. The iHMP IBD data set includes multiomics data from subjects with Crohn’s Disease (CD), ulcerative colitis (UC), and non-IBD (i.e., healthy), all of which were used in this study. MicrobiomeHD database includes 28 published case-control gut microbiome studies spanning ten diseases, from which we chose data sets associated with *C. difficile* infections and enteric diarrhea. These choices were made because the role of many taxa for those diseases are reasonably well established. Table 1 gives a summary of IBD related data sets. For each data set, we computed the total causal effect of each microbial taxon on all other taxa.

**Table 1.** Description of IBD-related data set

| Database | Data Set | # of Samples |
| --- | --- | --- |
| iHMP | Ulcerative colitis (UC) | 459 |
|  | Crohn’s disease (CD) | 749 |
|  | non-IBD (healthy) | 429 |
| Data (MicrobiomeHD) | Cases | Controls |
| EDD (Enteric diarrhea) | 201 | 82 |
| CDI (*C. difficile* infection) | 93 | 154 |

To explore the effect of antibiotics on antibiotic-resistant taxa, we analyzed a dataset from Gibson et al. [19]. It consists of 401 stool metagenomic samples from 84 premature infants that were sampled in multiple time points. All but two infants received antibiotic therapy within the first 24 hours. Sixty-one percent of the infants received additional antibiotic treatments (“Antibiotic” cohort) between 1–10 weeks of life. The remaining 39 percent formed the “Control” group. Each treatment consisted of one or more antibiotics. We considered each measurement as a separate sample from a distribution, and did not take into account the temporal aspect of the measurement as showed in [17].

### 3.3 Experiments

For each data set, we generated a causal structure by applying the PC-stable algorithm [10] and we computed the causal effect of each microbial taxon on all other taxa. We also computed the change in causal effects in healthy (non-IBD) versus diseased states.

To understand the causal role of microbial taxa in disease mechanisms we augmented the data set by merging healthy and disease abundance matrices and by adding an extra node named ‘disease’; we call this process ‘context embedding’. Context embedding is important for causal inference because in different contexts, the same event can be interpreted differently. For the healthy state, the value of disease node is 0, and for the disease state its value is 1. Thus the disease node becomes a binary random variable. We computed the causal effect of all taxa on disease, and vice versa.

For the antibiotic data set, as part of preprocessing, we profiled metagenomic reads against 14506 complete bacterial, archaeal, and viral genome sequences from RefSeq v.92 using FLINT [53] framework. Reference genomes were obtained from a repository hosted by the Kraken [58] tool. After obtaining abundance matrix, we created a causal network using recently used antibiotics and relative abundance of bacterial taxa. We computed causal effects of each antibiotic on each taxon and vice versa, to be used for further analysis.

## 4 Results and Analysis

### IBD and non-IBD Data

Fig. 1 shows a causal structure inferred from ulcerative colitis (UC) samples. Fig. 2 represents a causal network combining data from non-IBD and UC samples, but with an additional “disease” node (colored blue). Fig. 3 shows a causal network with nodes representing antibiotics and microbial taxa. Some more networks are shown and explained in the Appendix. In each network, nodes represent random variables for relative abundance of taxa, disease status, or antibiotic dosages. Edges represent their conditional relationships. The size of each node is proportional to sum of relative abundance and the color of the edges represents the sign of the correlation between the node variables. In a causal structure, directed edges suggest potentially direct causal effect between the connected variables. The absence of an edge suggests that there is no direct causal effect, although indirect causal effects may exist. An inferred causal structure may contain undirected edges if the data are not enough to support an edge orientation. Those undirected edges remain causally “uninterpretable”.

**Fig. 1.**
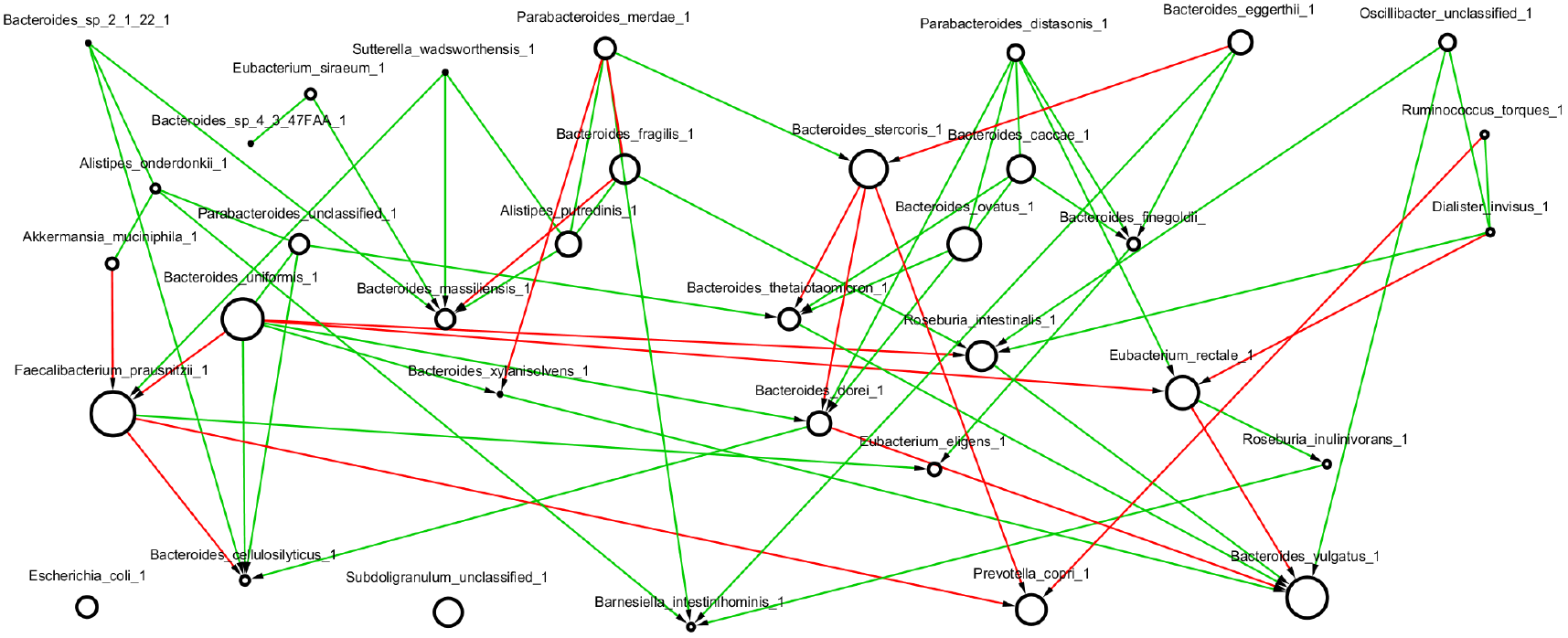
Causal network from Ulcerative Colitis (UC) data. All nodes represent taxa abundance.

**Fig. 2.**
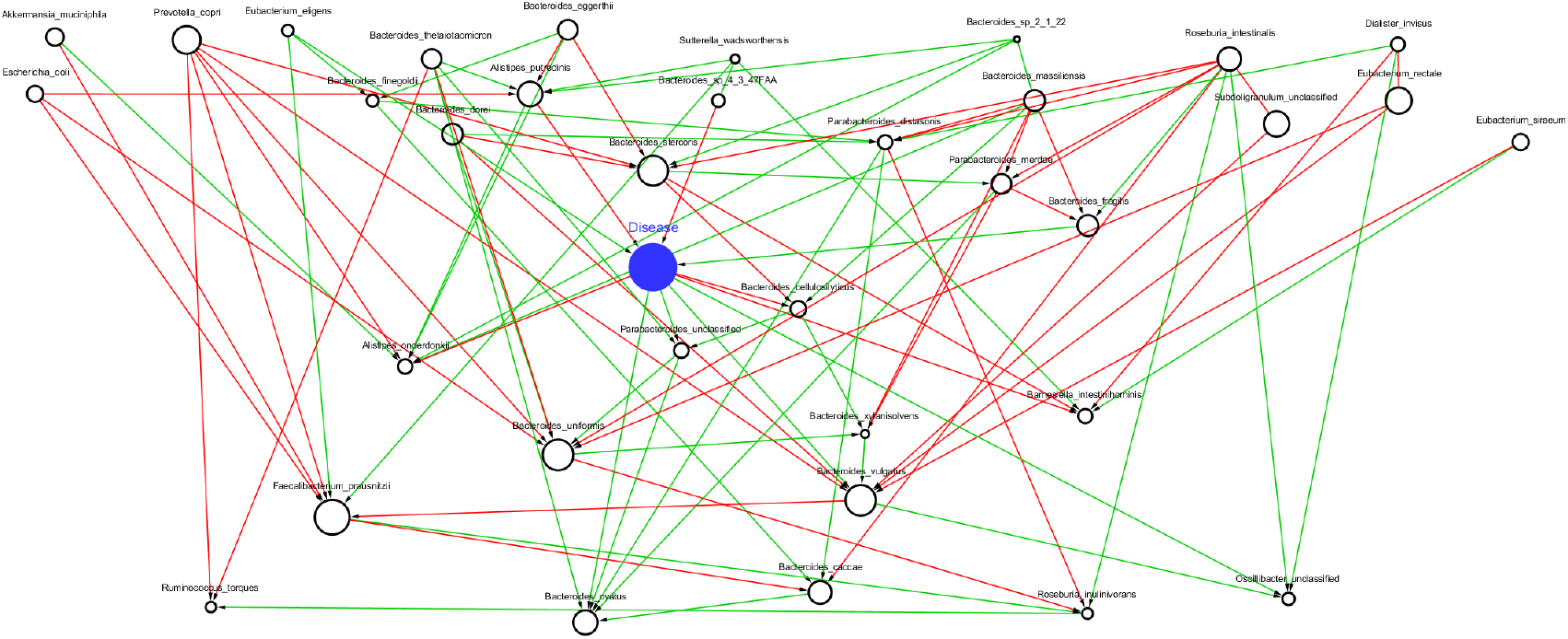
Causal network after introducing a “disease” node and using data from UC and non-IBD samples. Disease node is shown as a filled blue node.

**Fig. 3.**
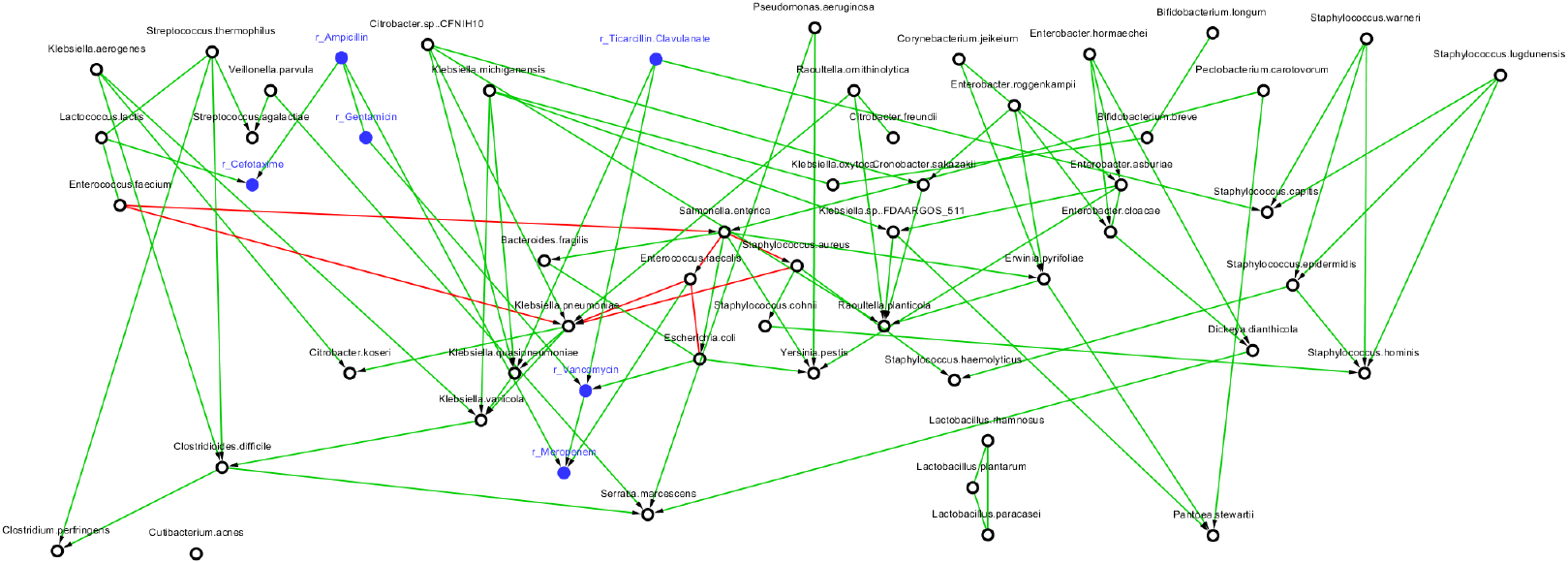
Causal network with antibiotics and taxa from samples obtained during or immediately after the antibiotic treatment. Blue nodes represent antibiotic doasges and black nodes represent taxa abundance.

We looked at the distribution of causal effects for each data set as shown in Fig. 4. Most of the causal effects are relatively small (see peak centered at 0). Approximately 30% of the causal effects are relatively large. The top 15% (shown in green rectangle) and bottom 15% (shown in red rectangle) are zoomed in for clarity. Based on sum of absolute values of causal effects we ranked the taxa as shown in Fig. 5. The hope is that this list shows the taxa that play a significantly key role in health and/or disease.

**Fig. 4.**
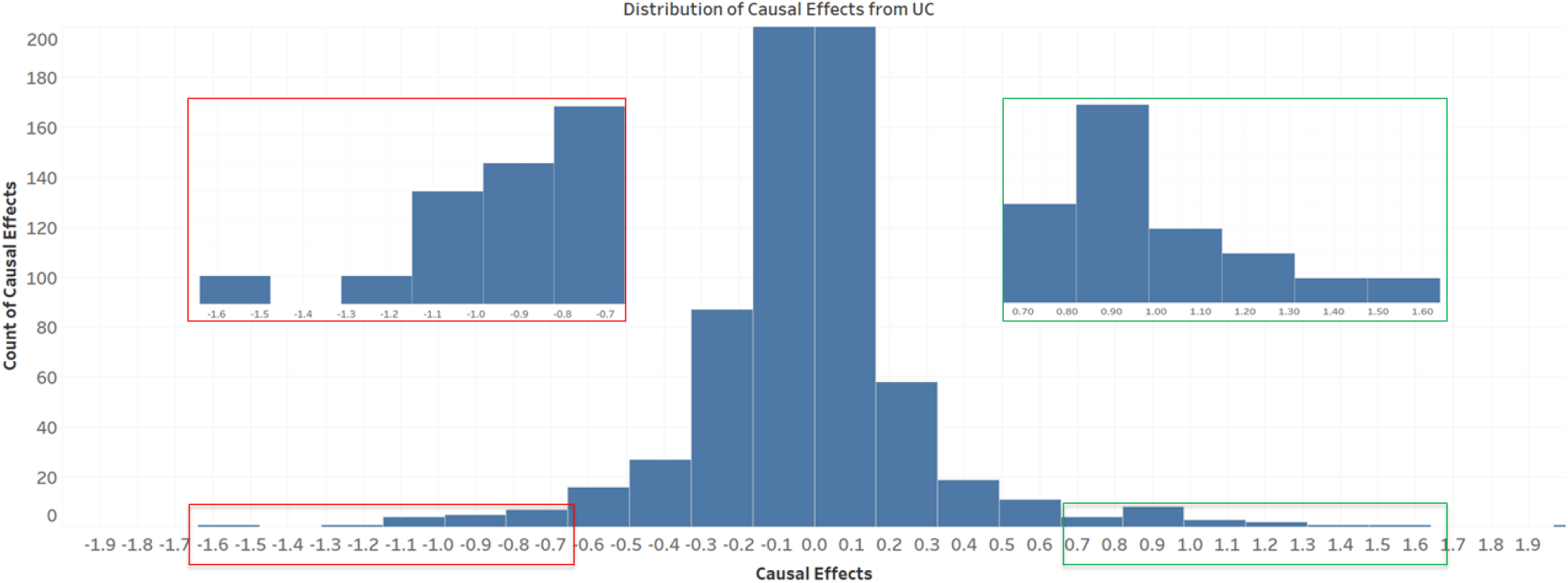
Histogram of causal effect values in UC. The top (green) and bottom (red) 15% are zoomed in for details.

**Fig. 5.**
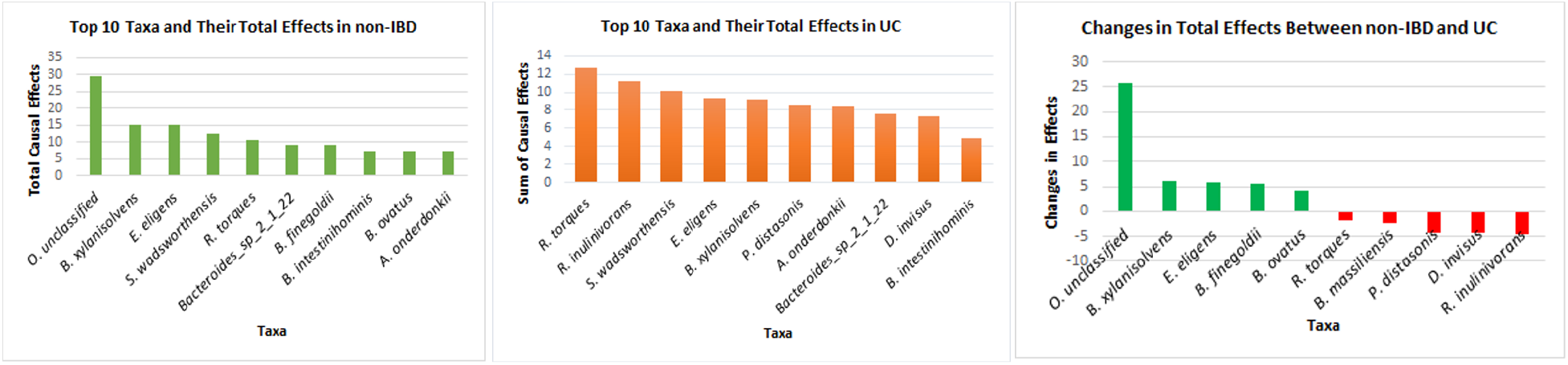
Top ten causally significant taxa from non-IBD (left), UC (middle), and Top 10 changes in causal effects from UC to non-IBD (right)

More interesting patterns are visible if we look at the changes in total causal effects between non-IBD (healthy) and UC taxa. Fig. 5 shows the ten taxa with the highest change in total causal effects. Green bars indicate higher total causal effect values in healthy, while red bars indicate higher values in UC, suggesting that the taxa on the left of the chart are potentially playing a beneficial role in health individuals while the taxa on the right of the chart are playing a harmful role in UC. Thus, in non-IBD subjects, the bacterial taxa *B. xylanisolvens*, *E. eligens*, *B. finegoldii*, *B. ovatus* and some species of *Oscillobacter* have more causal impact on the remaining taxa than others. These claims are supported by published literature, which show those taxa are potentially beneficial [28, 52, 32, 57]. On the other hand, in the diseased state (UC), other taxa including *R. torques*, *B. massiliensis*, *P. distasonis*, and *D. invisus* are more impactful. Again, the published literature supports the above claims [29, 26, 55, 33]. Thus our methods allow us to identify potentially beneficial and pathogenic bacteria in microbiomes.

### Disease networks

We measured causal effects of each taxa on the specially identified “disease node”. We sorted the absolute value of the effects as shown in Table 2. When we queried the published literature on this topic, we discovered that barring two, all the taxa listed in Table 2 are known to be either potentially pathogenic or beneficial, again supporting the claim that our approach helps to identify pathogenic and beneficial bacteria in healthy and diseased patients. We also computed total causal effects of disease on individual taxa. Most of the causal effects values are nearly zero, with values in the range of [−0.02, 0.02]. We then listed taxa with values greater than 0.01 or less than −0.01. Our findings suggest that the disease affects some commensal bacteria to become more abundant, while shrinking the abundance of potentially beneficial bacteria. This claim is supported by published literature as summarized in Table 3.

**Table 2.**
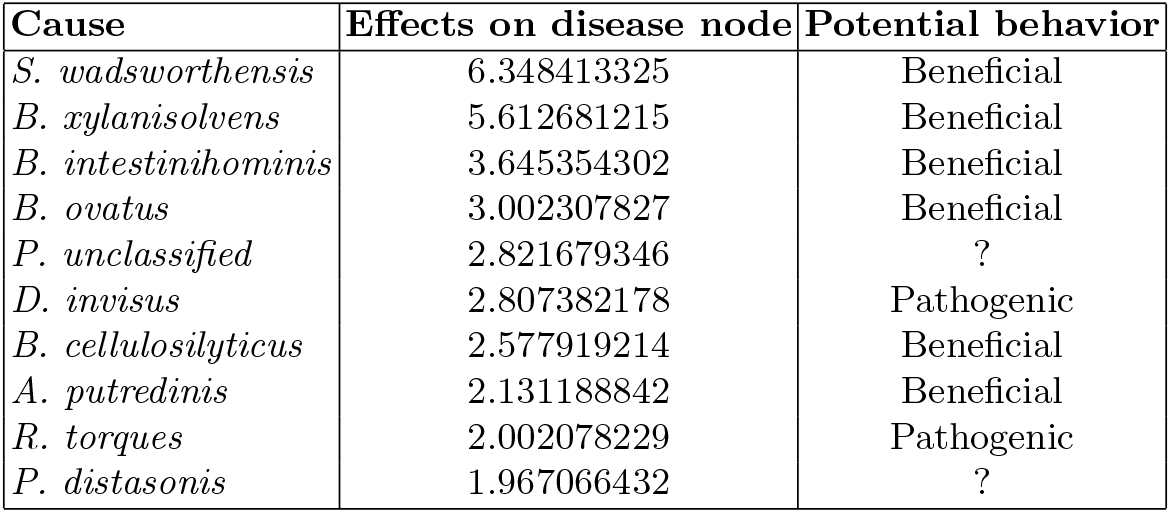
Taxa with highest causal effect on the *ulcerative colitis* “disease” node.

**Table 3.** Taxa most impacted by the *ulcerative colitis* “disease” node

| Taxa name | Behavior | References |
| --- | --- | --- |
| *B. uniformis* | Commensal | [43] |
| *B. ovatus* | Commensal | [44] |
| *E. coli* | Commensal | [50] |
| *A. muciniphila* | Beneficial | [61] |
| *P. copri* | Beneficial | [8] |
| *B. stercoris* | Beneficial | [31] |
| *A. putredinis* | Beneficial | [23] |

### Antibiotics Data

From the antibiotic data set, we learned a causal network and computed causal effects of each antibiotic to each taxa. The top seven positive causal effects and top seven negative causal effects are shown in Table 4 since all other values were close to the background noise. A positive causal effect of an antibiotic on a taxon suggests that either the antibiotic is inappropriate for that particular taxon or the taxon is resistant to the antibiotic. Similarly, a negative causal effect suggests that the antibiotic was effective against the taxon.

**Table 4.** Causal effects of antibiotics on taxa

| Cause | Effects | Magnitude of Causal Effects | Average $\Delta$ abundance |
| --- | --- | --- | --- |
| Ticarcillin Clavulanate | *K.pneumoniae* | 0.041646824 | + |
| Cefotaxime | *E.coli* | 0.037287043 | - |
| Meropenem | *E.faecalis* | 0.021974783 | + |
| Vancomycin | *E.coli* | 0.016519979 | 0 |
| Gentamicin | *E.coli* | 0.01563053 | + |
| Meropenem | *S.aureus* | 0.013329356 | NA |
| Ampicillin | *E.coli* | 0.010371501 | + |
| Meropenem | *K.pneumoniae* | -0.023983874 | - |
| Cefotaxime | *E.faecalis* | -0.017125561 | + |
| Cefotaxime | *K.pneumoniae* | -0.01689829 | - |
| Cefotaxime | *E.faecium* | -0.015538356 | - |
| Ticarcillin Clavulanate | *E.faecalis* | -0.009494291 | - |
| Vancomycin | *E.faecalis* | -0.009451166 | + |
| Ticarcillin Clavulanate | *S.aureus* | -0.009099242 | NA |

The significance of the causal graph shown in Figure 3 is that we are able to separate the effect of each antibiotic on a taxon. In general, doctors often administer combinations of antibiotics. In our study, 17 unique combinations of antibiotics were used to treat infants (one, two, or three types at a time). Therefore, it is difficult to identify which particular drug is associated with the rise of antibiotic-resistant taxa, or which antibiotic is most effective against pathogenic bacteria. Our proposed methods appear to be able to deconvolve these effects.

As mentioned above, the top 7 positive and negative causeal effect values of antibiotics on taxa are shown in Table 4. Except for three (out of 14) cases, our causal effect values can explain the average change in relative abundance that was computed by the previous study of Gibson et al. [19]. In particular, if the causal-effect of an antibiotic on taxa is positive, the average abundance of this taxa is increased and vice versa. The contradictory results may lead to new insights about antibiotic effectiveness. For example, even though after administering Cefotaxime and Vancomycin the relative abundance of *E. faecalis* on average tended to increase, our causal effect graph suggests that these antibiotics were effective against these taxa and that some other factors may be causing their increased abundance.

## 5 Conclusion

Causal inference shows promising results in analyzing microbiome data, especially in the identification of potentially pathogenic, beneficial, and antibiotic-resistant bacteria. Thus, in future, this process can allow us to evaluate the efficacy of probiotics and prebiotics. Moreover, causal inference from purely observational data is important to prioritize in picking wet-lab experiments for further anaysis. *Intervention* techniques can be used to quantify the average causal impact of one entity on another. The next challenge is to study the causal effect of one entity on another within a single sample.

